# Abscisic acid modulates auxin-responsive hypocotyl elongation

**DOI:** 10.1101/2021.04.25.441358

**Authors:** Ryan J. Emenecker, Joseph Cammarata, Irene Yuan, Lucia C. Strader

**Affiliations:** Department of Biology, Washington University, St. Louis, MO 63130, USA; Center for Science and Engineering Living Systems (CSELS), Washington University, St. Louis, MO 63130, USA; Center for Engineering Mechanobiology, Washington University, St. Louis, MO 63130, USA; Department of Biology, Duke University, Durham, NC 27708, USA

## Abstract

Auxin regulates many aspects of plant growth and development in concert with other plant hormones. Auxin interactions with these other phytohormones to regulate distinct processes is not fully understood. Using a forward genetics screen designed to identify seedlings resistant to the suppressive effects of auxin on dark-grown hypocotyl elongation, we identified a mutant defective in *ABA ALDEHYDE OXIDASE3* (*AAO3*), which encodes for the enzyme that carries out the final step in the biosynthesis of the plant hormone abscisic acid (ABA). We found that all examined ABA deficient mutants display resistance to the inhibitory effects of auxin on dark-grown hypocotyl elongation, suggesting that aspects of ABA signaling are downstream of auxin in regulating dark-grown hypocotyl elongation. Conversely, these mutants display wild type responsiveness to auxin in root elongation assays, suggesting that ABA does not act downstream of auxin in regulating elongation of the root. Our RNA-seq analysis suggests that many auxin-repressed genes in the hypocotyl require an intact ABA pathway for full repression. Our results suggest a model in which auxin partially requires intact ABA biosynthesis in order to regulate hypocotyl elongation, but not to regulate primary root elongation, suggesting that the genetic interactions between these two pathways are tissue-dependent.

## Introduction

Plant hormones act to coordinate plant growth and developmental processes (reviewed in 1). The plant hormone auxin is a central regulator of many plant growth and developmental processes through regulation of cell division and expansion (reviewed in 2). Because auxin is a central regulator of basic growth processes, it sits at the nexus of multiple signaling pathways, typically acting downstream in each of these signaling pathways. How auxin coordinates with other signalling pathways to regulate specific growth processes is not fully understood.

The plant hormone abscisic acid (ABA) is often thought of as a “stress hormone” although it also plays roles under non-stress conditions (3). Increasing evidence suggests that auxin and ABA work together to regulate various processes (reviewed in 4), including seed germination, (5-8), hypocotyl elongation (9), root elongation (6, 8, 10), lateral root formation (11-14), and cotyledon expansion (15). Notably, in all cases where it has been investigated, including seed germination, root elongation, and lateral root formation, disruption of either auxin biosynthesis or auxin signaling reduces the effects of ABA on growth processes (4).

Seedling establishment is a tenuous time in a plant’s life. After a seed germinates, typically below the soil line, the nascent seedling must quickly burn through seed reserves to fuel the growth necessary to emerge from the soil so that captured sunlight can become an energy source. In Arabidopsis, this growth of the hypocotyl is driven solely by cell expansion. Multiple pathways converge on regulating Arabidopsis hypocotyl elongation; however, auxin is a key regulator of the cell expansion driving this growth. Because many mutants that exhibit resistance to auxin in root growth assays are not markedly resistant to the inhibitory effects of auxin on dark-grown hypocotyl elongation (10, 16), we hypothesized that our understanding of how auxin regulates this critical process was incomplete.

Here, we describe the identification of a mutant resistant to the suppressive effects of supplied auxin on dark-grown hypocotyl elongation. The *ABA ALDEHYDE OXIDASE3* (*AAO3*) gene is required for full responsiveness to auxin in the regulation of hypocotyl elongation. AAO3 is the enzyme responsible for catalyzing the final step in ABA biosynthesis (17). Examination of various ABA biosynthetic and signaling mutants revealed that auxin requires functional ABA biosynthesis and signaling to fully regulate hypocotyl elongation, suggesting that aspects of ABA function downstream of auxin in this tissue. We found that the root and the shoot have distinct transcriptional responses to auxin and that many auxin-responsive transcripts in the hypocotyl, but not the root, require intact ABA biosynthesis. In particular, the repression of genes by auxin was limited by defects in ABA biosynthesis. Our data support a model in which auxin relies on intact ABA biosynthesis to regulate some aspects of hypocotyl elongation. Our findings further suggest that some tissues can display ABA-auxin interactions distinct from other tissues and raise the possibility that ABA may alter auxin-regulated hypocotyl growth under stressful conditions.

## Results

### Mutations in *ABA ALDEHYDE OXIDASE3* confer auxin resistance in dark-grown hypocotyl elongation assays

Depending on context, auxin may either promote or inhibit cell expansion (reviewed in 2). Exogenous auxin inhibits hypocotyl elongation in dark-grown seedlings (16); however, many mutants strongly resistant to the inhibitory effects of auxin in root elongation assays are only mildly resistant to the inhibitory effects of auxin on dark-grown hypocotyl elongation, consistent with the possibility that distinct mechanisms drive auxin responses in hypocotyl and root tissues. We therefore expanded from our previous hypocotyl resistance (HR) screen (16) to identify ethyl methanesulfonate (EMS)-mutagenized dark-grown M_2_ seedlings that display resistance to the inhibitory effects of the auxin indole-3-butyric acid (IBA) on hypocotyl elongation. Hypocotyl Resistant 12 (HR12) displayed resistance to the inhibitory effects of the natural active auxin indole-3-acetic acid (IAA), the natural long-chain auxin precursor IBA, the synthetic active auxin 2,4-dichlorophenoxyacetic acid (2,4-D), and the synthetic long-chain auxin precursor 2,4-dichlorophenoxybutyric acid (2,4-DB) on dark-grown hypocotyl elongation (Figures 1A, 1B). This data suggests that the defect in HR12 generally affects auxin responsiveness in the hypocotyl. Although HR12 displayed resistance to the inhibitory effects of auxin on dark-grown hypocotyl elongation (Figures 1A, 1B), HR12 displayed wild-type sensitivity to the inhibitory effects of auxins on light-grown root elongation (Figure 1C). Thus, HR12 appears to be specifically defective in response to auxin in hypocotyl elongation assays.

**Figure 1.**
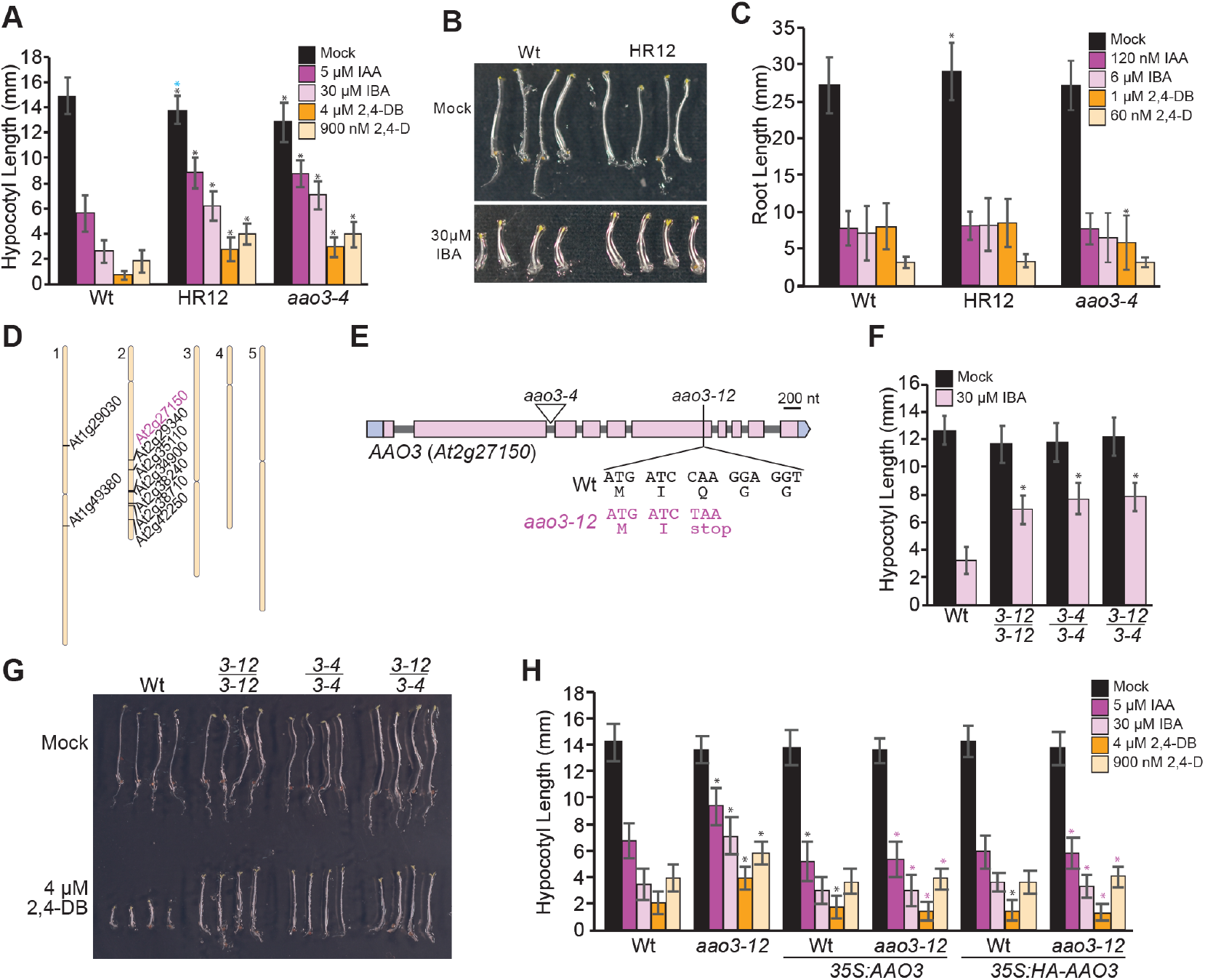
Characterization of HR12 and confirmation of the causative mutation. (A) Mean hypocotyl lengths (± SD; *n* ≥ 78) of wild type (Wt; Col-0), HR12, and *aao3-4* seedlings grown in the dark on media supplemented with the indicated treatment. Statistically significant (*P* ≤ 0.01) differences from wild type are indicated with an asterisk and significant differences (*P* ≤ 0.01) in comparison with *aao3-12* are marked with a purple asterisk. (B) Photograph of wild type (Wt; Col-0) and HR12 seedlings grown in the dark on media supplemented with ethanol (mock) or 30 µM IBA. (C) Mean root lengths (± SD; *n* = 80) of wild type (Wt; Col-0), HR12, and *aao3-4* grown under continuous illumination on media supplemented with the indicated treatment. Significant differences (*P* ≤ 0.01) from wild type are indicated with an asterisk. (D) Bulk segregant analysis revealed homozygous EMS mutations within the HR12 backcrossed population. (E) *At2g27150*/*AAO3* gene schematic depicting the EMS-consistent point mutation identified in *aao3-12* (HR12) and the *aao3-4* T-DNA insertion allele. *aao3-12* (HR12) carries a C-to-T mutation at position 3846 (where the A of ATG is +1) that results in a Q1081-to-stop substitution. (F) Non-complementation assay of HR12 (*aao3-12*) and *aao3-4* (SALK-072361c). Mean hypocotyl lengths (± SD; *n* = 20) of dark-grown seedlings for the indicated genotype for Mock and 30 ⌈M IBA treatments. Significant differences (*P* ≤ 0.01) between wild type and the indicated genotype are indicated by a black asterisk. (G) Photograph of seedlings grown in the dark on media supplemented with ethanol (mock) or 4 ⌈M 2,4-DB treatment. Examined genotypes include wild type (Wt; Col-0), *aao3-12, aao3-4*, and the F_1_ trans-heterozygote from a cross between *aao3-12* and *aao3-4 (aao3-12/aao3-4)*. (H) The *AAO3* gene complements *aao3-12* (HR12) phenotypes. Mean hypocotyl lengths (± SD; *n* = 40) of wild type (Wt) and *aao3-12* seedlings carrying no transgene, *35S:AAO3*, or *35S:HA-AAO3* grown in the dark on media supplemented with mock (ethanol) or the indicated auxin. Significant differences (*P* ≤ 0.01) in comparison with wild type are indicated with a black asterisk and significant differences (*P* ≤ 0.01) in comparison with *aao3-12* are marked with a purple asterisk.

To identify the causative mutation in HR12, we backcrossed the mutant to wild type and isolated auxin-resistant individuals in the F_2_ generation. We then used a whole genome sequencing of bulk segregants approach (6, 18) on DNA from pooled F_3_ individuals to identify nine homozygous EMS-induced mutations in the HR12 mutant (Figure 1D). One of these mutations was in *At2g27150* and resulted in a premature stop codon in *ABA ALDEHYDE OXIDASE3* (*AAO3*), prompting us to further investigate whether this mutation was causative for the auxin resistance seen in HR12. We named the *AAO3* mutation in HR12 *aao3-12* (Figure 1E).

To determine whether the *aao3-12* mutation was causative for the auxin resistance observed in HR12, we examined auxin responsiveness of a T-DNA insertional allele, Salk_072361C (hereafter referred to as *aao3-4*; Figure 1E). Similar to HR12, the *aao3-4* mutant displayed resistance to auxin in dark-grown hypocotyl elongation assays (Figure 1A) and displayed wild-type sensitivity to auxin in root elongation assays (Figure 1C), suggesting that *AAO3* defects may be causative for the observed auxin resistance in hypocotyl elongation assays in HR12. It is important to note that the *aao3-4* allele showed a very slight but statistically significant difference from wild type in the root elongation assays when treated with the synthetic auxin 2,4-DB. However, because we did not see statistically significant differences between wild type and *aao3-4* for any of the other treatments, and none of the differences between *aao3-12* and wild-type were significant, this small difference between wild-type and *aao3-4* when treated with 2,4-DB while statistically significant is unlikely to be biologically relevant.

To further confirm whether the *AAO3* mutation was causative in HR12, we crossed *aao3-4* to HR12 and examined auxin sensitivity of the F_1_ progeny, finding that the *aao3-12*/*aao3-4* trans-heterozygote displayed resistance to the inhibitory effects of IBA (Figure 1F) and 2,4-DB (Figure 1G) on hypocotyl elongation. Thus, *AAO3* defects are causative for the auxin resistance displayed in these mutants. As a final means of verifying that the *aao3-12* lesion caused the observed resistance to the inhibitory effects of auxin in dark-grown hypocotyl elongation, we transformed *aao3-12* with a wild type, genomic copy of AAO3 under the *35S* promoter. Expression of a wild-type copy of *AAO3* was sufficient to restore auxin sensitivity to *aao3-12* (Figure 1H). We should note that in the hypocotyl elongation assays using either IAA or 2,4-DB, wild type lines expressing *AAO3* under the *35S* promoter displayed increased sensitivity to auxin treatment in comparison to untransformed wild type (Figure 1H). While statistically significant, this difference was very small and not consistent across all auxins tested. Nonetheless, taken together, these results further support that the auxin response defects in HR12/*aao3-12* resulted from reduced AAO3 function.

### Additional ABA mutants exhibit auxin resistance in dark-grown hypocotyl elongation assays but not root elongation assays

AAO3 catalyzes the final step in ABA biosynthesis (Fig 2A) (17). We reasoned that if disruption of ABA biosynthesis, rather than an unrelated role for AAO3, causes the auxin resistance seen in *aao3* mutants, then mutants with defects in other steps in the ABA biosynthetic pathway should also display auxin resistance in dark-grown hypocotyl elongation assays. We therefore obtained T-DNA insertional alleles for *ABA DEFICIENT2 (ABA2)* and *ABA DEFICIENT3 (ABA3)*. We found that disruption of either ABA2 or ABA3 resulted in auxin resistance in dark-grown hypocotyl elongation assays, supporting a model where ABA biosynthesis is necessary for full auxin-mediated inhibition of dark-grown hypocotyl elongation (Figure 2B). In addition, similar to the *aao3* mutants, mutants defective in *ABA2* and *ABA3* displayed wild type sensitivity to auxin in root elongation assays (Figure 2C). This finding that ABA biosynthesis was required for auxin-mediated dark-grown hypocotyl elongation is in contrast with previous research that found that ABA is not necessary for auxin-induced inhibition of root elongation (6). However, our data clearly show that under the conditions tested here, auxin requires intact ABA biosynthesis in order to exert its full effects on hypocotyl elongation. Importantly, whereas the ABA biosynthesis mutants we examined dispaly resistance to auxin in the inhibition of dark-grown hypocotyl elongation, the ABA biosynthesis mutants are still capable of responding to auxin treatment in these assays; the response of the ABA mutants to auxin’s effects on hypocotyl elongation are attenuated compared to wild type. The requirement of intact ABA biosynthesis for full auxin-mediated inhibition of hypocotyl elongation suggests that some, but not all, aspects of ABA are downstream of auxin in the regulation of this process. Because ABA-deficient mutants are still moderately responsive to auxin, aspects of auxin-mediated regulation of hypocotyl elongation are clearly independent of ABA.

**Figure 2.**
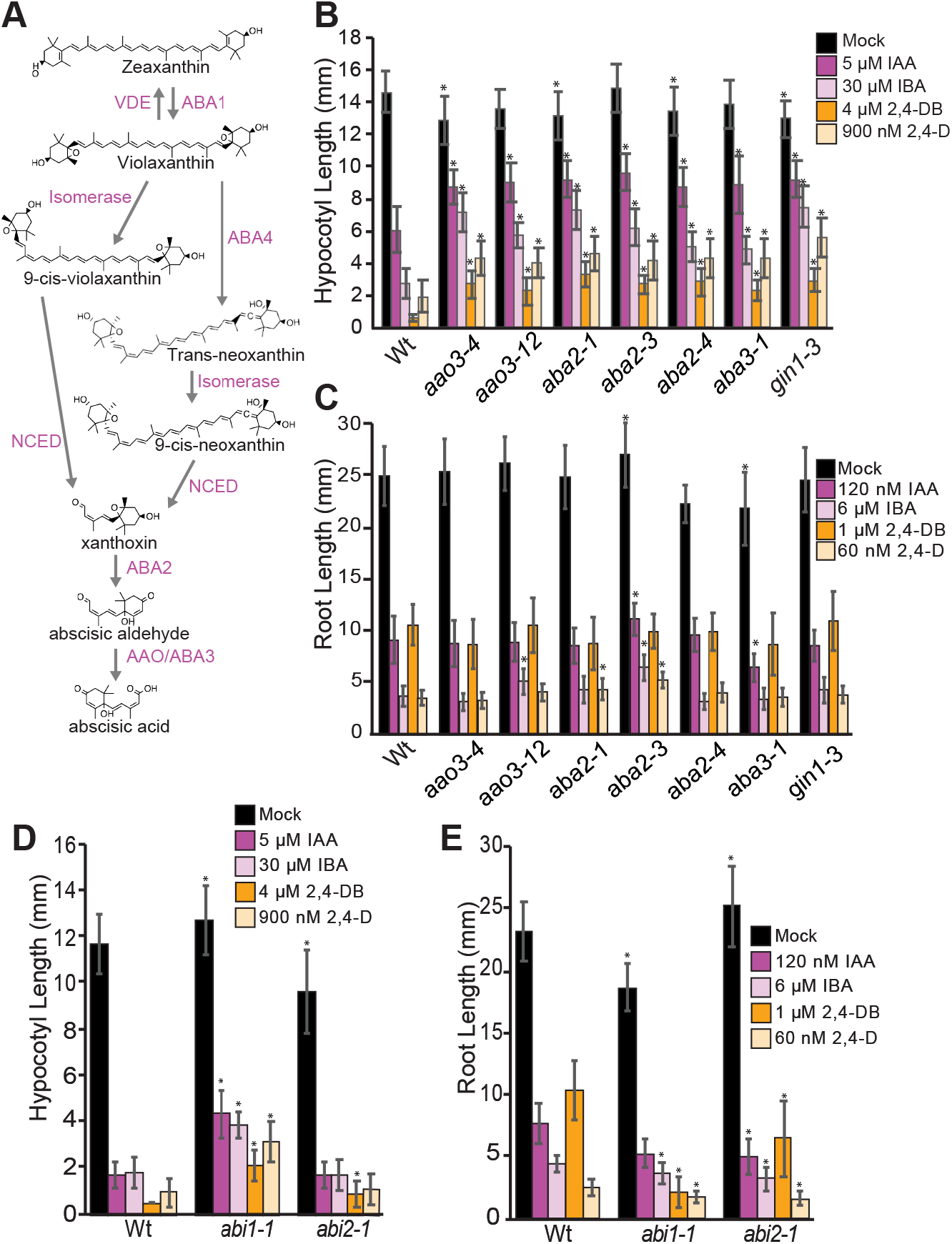
Assessing the role of ABA biosynthesis and signaling in hypocotyl and root elongation. (A) Schematic of the ABA biosynthetic pathway with intermediates and enzymatic steps indicated. (B) Mean hypocotyl lengths (± SD; *n* = 40) of wild type (Wt; Col-0), *aao3-4, aao3-12, aba2-1, aba2-3, aba2-4, aba3-1*, and *gin1-3* seedlings grown in the dark on media supplemented with ethanol (Mock) or the indicated hormone. The *gin1-*3 mutant is defective in *ABA2*. Significant differences (*P* ≤ 0.01) in comparison with wild type are marked by an asterisk. (C) Mean root lengths (± SD; *n* ≥ 35) of wild type (Wt; Col-0), *aao3-4, aao3-12, aba2-1, aba2-3, aba2-4, aba3-1*, and *gin1-3* seedlings grown in the light on media supplemented with ethanol (Mock) or the indicated hormone. Significant differences (*P* ≤ 0.01) in comparison to wild type are indicated by an asterisk. (D) Mean hypocotyl lengths (± SD; *n* = 40) of wild type (Wt; L*er*-0), *abi1-1*, and *abi2-1* seedlings grown in the dark on media supplemented with ethanol (Mock) or the indicated hormone. Significant differences (*P* ≤ 0.01) in comparison to wild type are indicated by an asterisk. (E) Mean root lengths (± SD; *n* ≥ 32) of wild type (Wt; L*er*-0), *abi1-1*, and *abi2-1* seedlings grown in the light on media supplemented with ethanol (Mock) or the indicated hormone. Significant differences (*P* ≤ 0.01) in comparison to wild type are indicated by an asterisk.

Binding of ABA to the PYROBACTIN RESISTANCE/PYROBACTIN 1-LIKE/REGULATORY COMPONENTS OF ABA RECEPTOR (PYR/PYL/RCAR) family of proteins results in inhibition of the clade A members of the PROTEIN PHOSPHATASE 2C (PP2C) family of protein phosphatases (19, 20). The clade A PP2C phosphatases typically inhibit the ABA response, and their inhibition by PYR/PYL/RCAR family members in the presence of ABA results in reactivation of other ABA signaling components including the SLOW ANION CHANNEL ASSOCIATED1 (SLAC1) ion channel and the SUCROSE NONFERMENTING1-RELATED PROTEIN KINASES2 (SnRK2) kinases (21, 22). To further investigate the role of ABA in auxin-mediated regulation of hypocotyl elongation, we examined two gain-of-function ABA signaling mutants, *aba insensitive1-1* (*abi1-1*) and *aba insensitive2-1* (*abi2-1*), both of which display strong resistance to ABA. ABI1 and ABI2 are in the PP2C family of protein phosphatases; the *abi1-1* and *abi2-1* gain-of-function mutants display strong resistance to ABA. We found *abi1-1* but not *abi2-1* displayed resistance to all four auxins tested in dark-grown hypocotyl elongation assays (Figure 2D). This is consistent with a previous study that found dephosphorylation of plasma membrane H+ATPases by ABA, which has been proposed to be a possible mechanism by which ABA can inhibit hypocotyl elongation, is diminished in the *abi1-1* mutant (23). In contrast to a previous study in which *abi1-1* and *abi2-1* displayed wild type sensitivity to auxin in root elongation assays (6), we found that both *abi1-1* and *abi2-1* exhibited a slight hypersensitivity to auxin in these assays (Figure 2E). These results may suggest that aspects of ABA signaling are involved in auxin-mediated regulation of root elongation; however, more extensive characterization would be required to draw this conclusion. These findings support that ABA biosynthesis and some aspects of ABA signaling are necessary for auxin to fully exert its inhibitory effects on dark-grown hypocotyl elongation, but auxin-mediated regulation of root elongation does not require intact ABA biosynthesis.

### ABA mutants are resistant to auxin in hypocotyls independent of light conditions

Our described root elongation assays were carried out under light-grown conditions whereas our hypocotyl elongation assays were carried out under dark-grown conditions, raising the possibility that differences between the root elongation and hypocotyl elongation assays in dependence on ABA biosynthesis for auxin responsiveness may be a light-dependent phenomenon rather than specific to the tissues examined. To test this possibility, we carried out hypocotyl elongation assays under continuous exposure to light. In light-growth conditions, supplied auxin promotes hypocotyl elongation (24). For light-grown hypocotyl elongation assays, we found that the ABA biosynthesis mutants were resistant to the synthetic auxin picloram in promotion of hypocotyl elongation (Figure 3A, Figure 3B), although the ABA biosynthesis mutants displayed longer hypocotyls than wild type in the control treatment (Figure 3A). The lengthened hypocotyls of the ABA biosynthesis mutants under mock conditions suggests endogenous ABA may function to reduce hypocotyl elongation under light-grown conditions. Furthermore, the reduced percent increase in hypocotyl length seen in the ABA biosynthesis mutants in response to auxin may suggest that auxin is less able to promote hypocotyl elongation in the ABA biosynthesis mutants as compared to wild type. This would support the notion that aspects of auxin-mediated regulation of hypocotyl elongation depend on intact ABA biosynthesis independent of whether the seedlings are grown in the dark or in the light. However, because the ABA biosynthesis mutants had longer hypocotyls under mock conditions, it is also possible that the ABA biosynthesis mutants did not undergo further hypocotyl elongation in response to picloram because they are maximally elongated even in the absence of auxin.

**Figure 3.**
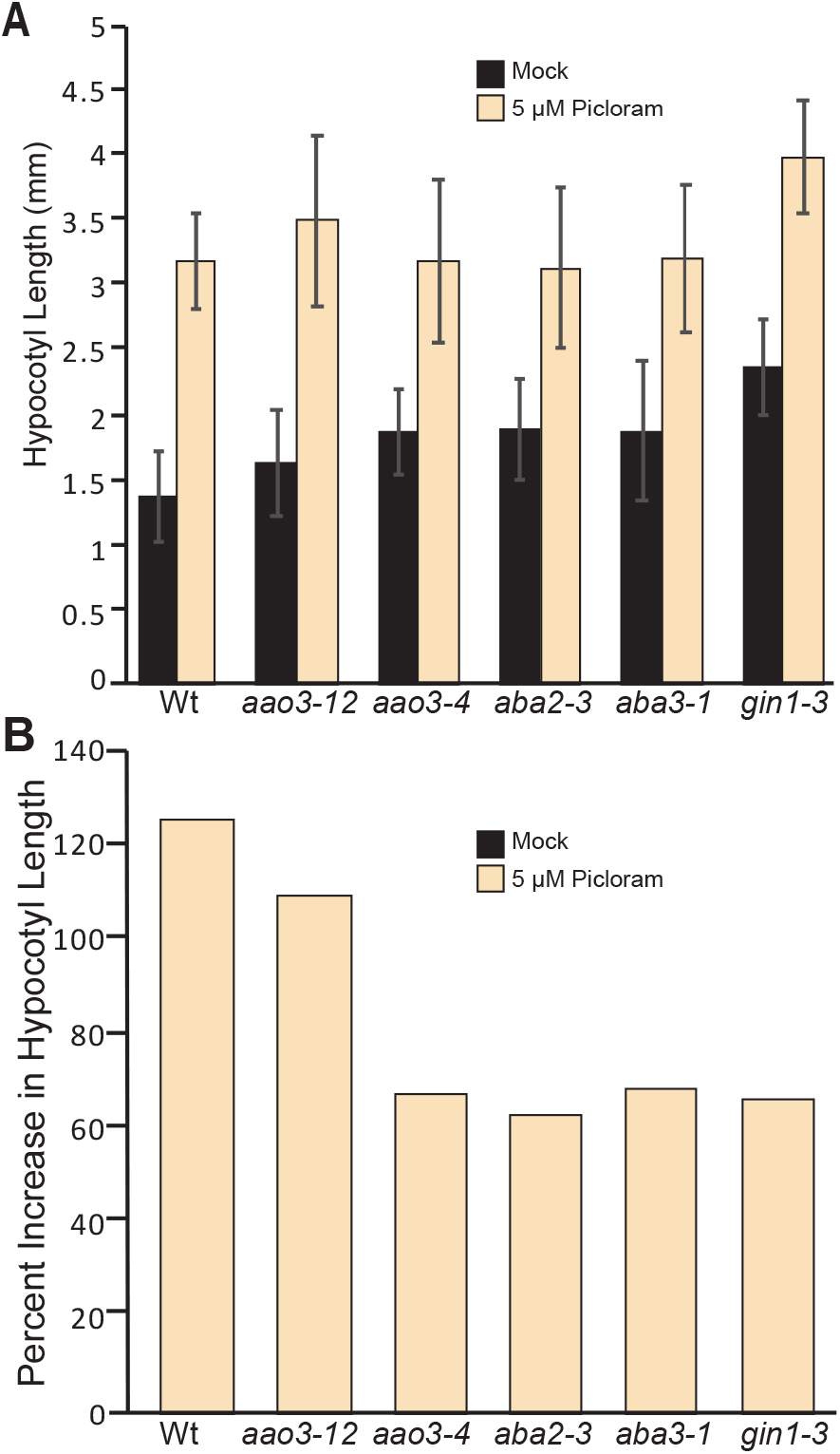
ABA biosynthesis mutants are resistant to promotion of hypocotyl elongation by auxin in the light. (A) Mean hypocotyl lengths (± SD; *n* = 30) of wild type (Wt; Col-0), *aao3-12, aao3-4, aba2-3, aba3-1*, and *gin1-3* seedlings grown under continuous illumination on media supplemented with ethanol (Mock) or the synthetic auxin picloram. (B) Mean percent increase in hypocotyl lengths of seedlings grown in the presence of 5 ⌈M picloram compared to those grown on media supplemented with ethanol (Mock) using the data from Figure 3A.

### Auxin and ABA show largely distinct impacts on transcript accumulation in the hypocotyl and the root

Our hormone response data suggest that auxin requires intact ABA biosynthesis to fully regulate hypocotyl elongation whereas auxin does not require intact ABA biosynthesis to regulate root elongation. To further shed light on the differences in ABA-dependent, auxin-mediated regulation of these two tissues, we examined the auxin- and ABA-regulated transcriptional programs in dark grown roots (everything below the root-hypocotyl junction) and dark-grown shoots (everything above the root-hypocotyl junction) (Figure 4A). Working under green safe lights, we treated four-day old dark-grown wild type seedlings with mock, IAA, or ABA for 2 hours prior to bisecting the root and the shoot at the root-hypocotyl junction and examining the differential effects of these hormone treatments in these two tissues. We found that auxin regulates largely non-overlapping sets of genes in dark-grown root and shoot tissues (Figure 4B), consistent with a model in which auxin has distinct effects in these two tissues. Through gene ontology analysis, we found that differentially expressed genes in the root included those with annotated roles in cell wall loosening, lateral root formation/root system development, steroid biosynthesis, and gravitropism (25). In contrast, genes that were differentially expressed in response to auxin treatment in the shoot included genes with roles in shade avoidance, positive regulation of growth, and response to brassinosteroid (25). Gene ontology analysis (25) revealed enrichment of auxin, auxin transport, regulation of transcription, and auxin homeostasis genes that differentially accumulate in response to auxin in both the root and the shoot.

**Figure 4.**
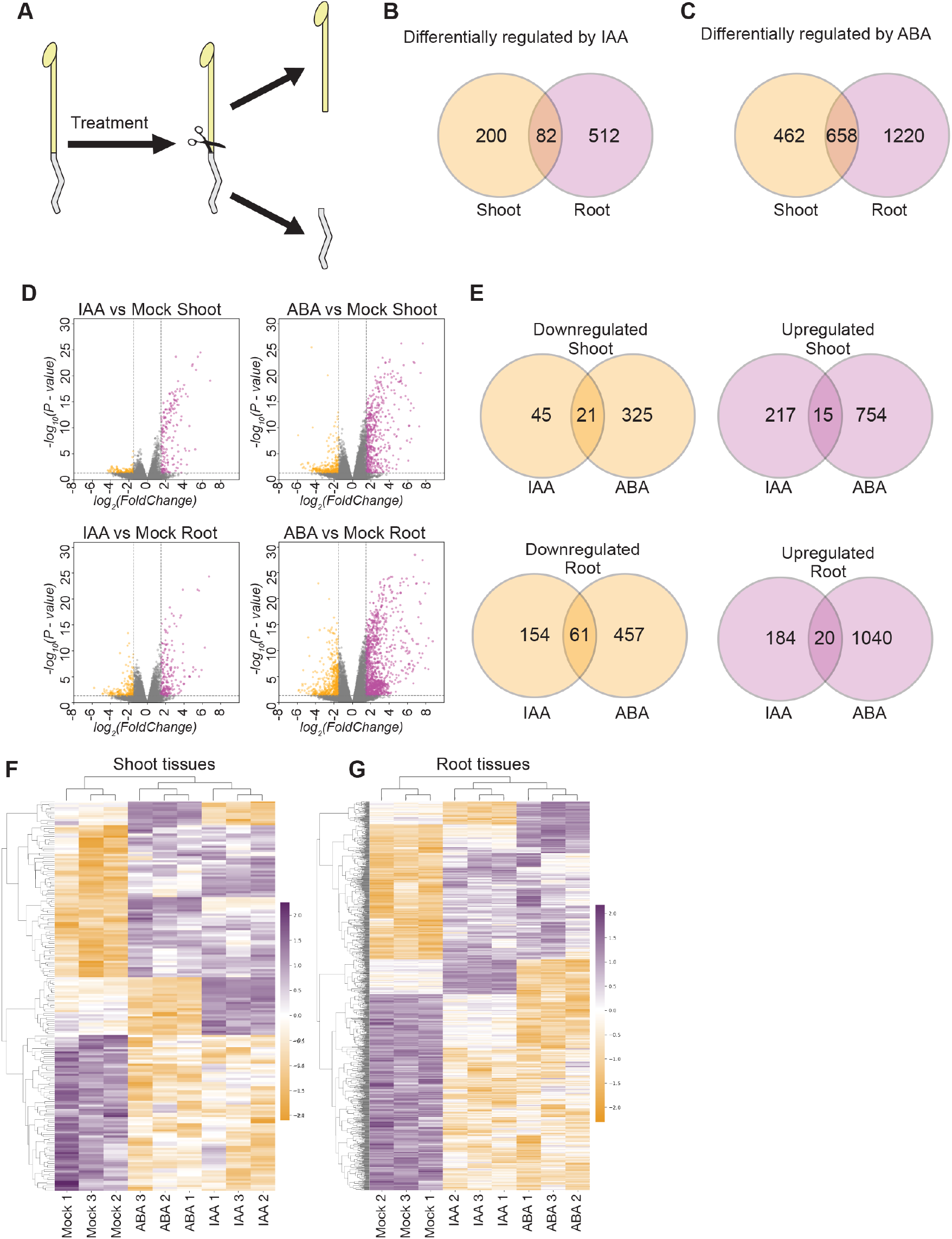
Auxin and ABA have distinct impacts on the transcription of genes in the root and the shoot. (A) Dark-grown seedlings were treated with ethanol (Mock), 10 ⌈M IAA, or 10 ⌈M ABA prior to dissection of the root and the shoot by cutting at the root-hypocotyl junction. RNA was extracted from the separated tissues, which was then used for RNA-seq analysis. (B) Venn diagrams showing the overlap between the datasets of genes differentially expressed (log fold change ≥ |1.0|; FDR ≤ 0.05) in response to auxin treatment the root and the shoot of wild type (Wt; Col-0). (C) Venn diagrams showing the overlap between the datasets of genes differentially expressed (log fold change ≥ |1.0|; FDR ≤ 0.05) in response to ABA treatment the root and the shoot of wild type (Wt; Col-0). (D) Volcano plots displaying pairwise transcript accumulation differences between auxin or ABA and mock-treated wild type (Col-0) shoots (top) and roots (bottom). (E) Venn diagrams showing the overlap between the datasets of genes differentially expressed (log fold change ≥ |1.0|; FDR ≤ 0.05) in response to auxin or ABA treatment in shoot or root tissues. (F) Hierarchical clustering of genes displaying differential expression among hormone-treated shoot samples (FDR ≤ 0.05). (G) Hierarchical clustering of genes displaying differential expression among hormone-treated root samples (FDR ≤ 0.05).

In contrast to previously published transcriptome profiles of auxin- and ABA-regulated genes (24, 26-28), our dataset was collected using dark-grown seedlings. Although it is difficult to compare across experiments, we found fewer auxin- and ABA-responsive genes in dark-grown seedlings than we expected based on previous experiments in light-grown seedlings. Future work would be necessary to distinguish whether these differences are truly reflective of differences in the biology between seedlings grown under these two light regimes.

We found little overlap in auxin- and ABA-responsive genes in either the root or the shoot of dark-grown seedlings (Figure 4E,F,G). This data is in contrast to our growth response assays (Figures 1, 2, 3) that suggest that ABA acts downstream in regulating dark-grown hypocotyl elongation. However, given that these two hormones regulate genes that have impacts other than simply regulating the elongation of the root or the hypocotyl, it is possible that the genes that have opposing effects in response to auxin and ABA treatment may be involved in other processes. In examining the15 genes that were upregulated by both ABA and IAA in the shoot, five were identified as genes that have previously been annotated as having roles in regulating hypocotyl elongation including SHI-RELATED SEQUENCE 5 (AT1G75520) (29), INDOLE-3-ACETIC ACID CARBOXYL METHYLTRANSFERASE (AT5G55250) (30), CYCLING DOF FACTOR 5 (AT1G69570) (31), SHORT HYPOCOTYL 1 (At1G52830) (32), OVATE FAMILY PROTEIN 1 (AT5G01840) (33). No genes that were downregulated by both IAA and ABA in the shoot have previously described roles in hypocotyl elongation.

### The *aba2* mutant reveals ABA-dependent auxin-regulated gene transcription

Our growth response data suggest that ABA biosynthesis is required for the full inhibitory effects of auxin on dark-grown hypocotyl elongation, but our RNASeq data suggest that auxin and ABA elicit distinct transcriptional responses in dark-grown hypocotyl tissue. We therefore sought to determine whether aspects of auxin transcriptional response are affected when ABA biosynthesis is disrupted.

Full auxin responses in hypocotyl-based growth assays require intact ABA biosynthesis. The 9-CIS-EPOXYCAROTENOID DIOXYGENASE (NCED) family regulates first committed step of ABA biosynthesis and is often transcriptionally regulated to affect ABA levels (34). Examination of *NCED* transcript accumulation revealed that *NCED5* transcripts were upregulated by auxin treatment in shoot tissues (Figure 5A), consistent with the possibility that auxin treatment upregulates ABA levels in this tissue.

**Figure 5.**
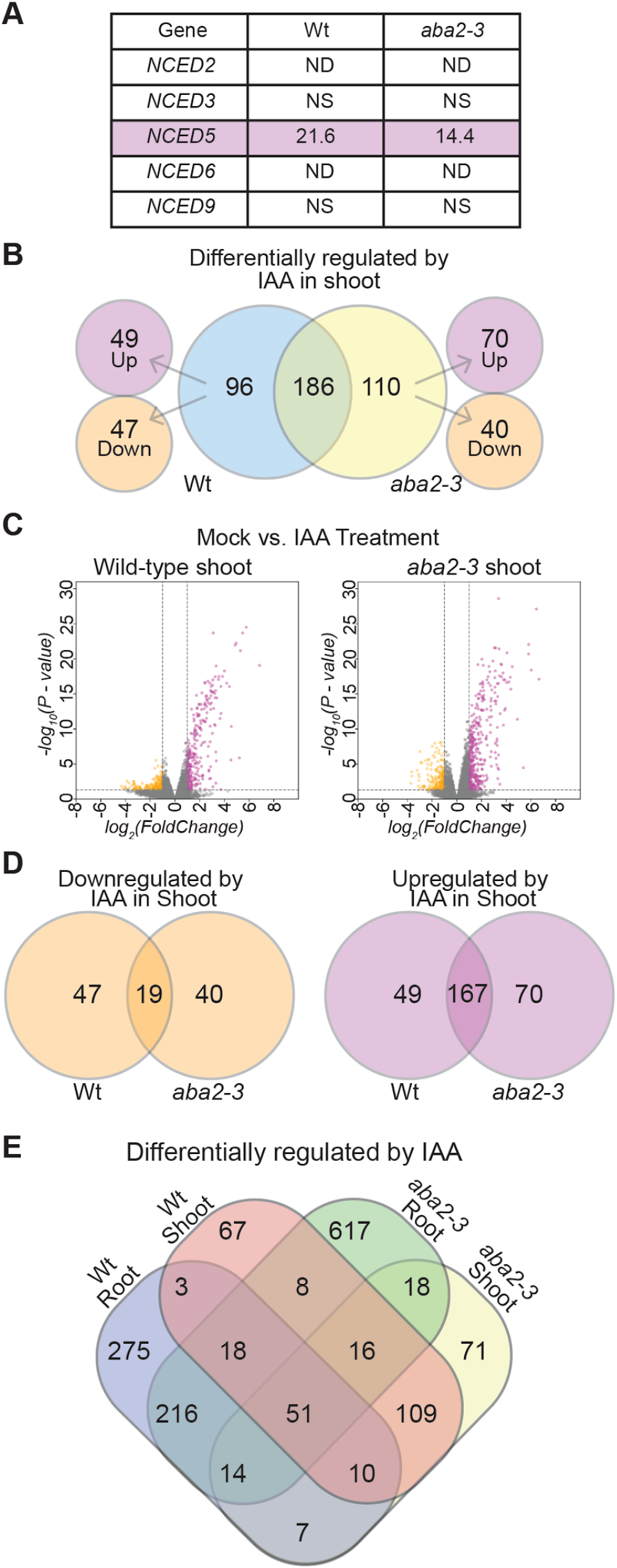
Disruption of ABA biosynthesis impacts the transcriptional response to auxin in the root and the shoot. (A) Auxin-responsive fold-change in *NCED* family member transcript accumulation in the shoots of wild type (Wt; Col-0) and *aba2-3* seedlings. ND designates not detected in RNA-seq experiment, (NS) designates not statistically significant (FDR ≤ 0.05). (B) Venn diagram showing overlap between the datasets of genes that are differentially expressed (log fold change ≥ |1.0|; FDR ≤ 0.05) in response to auxin treatment in the shoots of *aba2-3* and wild type (Wt; Col-0). (C) Volcano plots displaying pairwise transcript accumulation differences between mock and auxin-treated wild type (Wt; Col-0) (left) and *aba2-3* (right) shoot tissues. (D) Venn diagrams showing overlap between the datasets of genes that display decreased (left) or increased (right) accumulation (log fold change ≥ |1.0|; FDR ≤ 0.05) in response to auxin treatment of wild type (Wt; Col-0) or *aba2-3* shoot tissues. (E) Venn diagram showing overlap between the datasets of genes that are differentially expressed (log fold change ≥ |1.0|; FDR ≤ 0.05) in response to auxin treatment in the shoots of wild type (Wt; Col-0) and *aba2-3* and wild type.

Because *aba2* is defective in ABA biosynthesis (35), we reasoned that comparing the auxin-induced transcriptional response of *aba2* to wild type in the hypocotyl would allow us to identify auxin regulated transcripts that depend on intact ABA biosynthesis. Notably, ABA2 is one step downstream of NCED5 (Figure 2A); therefore, in the *aba2-3* mutant we would not expect upregulation of NCED5 to result in altered ABA levels. We found striking differences in auxin-regulated transcripts between wild type and *aba2* in dark-grown shoot tissues (Figure 5B, 5C). Further, transcripts downregulated by auxin were more strongly affected by loss of endogenous ABA than those upregulated by auxin (Figure 5D). To determine possible ABA biosynthesis-dependent mechanisms through which auxin regulates hypocotyl elongation, we identified 67 auxin-responsive genes that required ABA2 and were differentially regulated specifically in the shoot of wild type but not of *aba2* (Figure 5E, Supplemental File 1). We also identified 71 genes that were differentially regulated by auxin treatment in the shoot of *aba2* but not in the shoot of wild type (Figure 5E, Supplemental File 2). These data suggest that not only does auxin rely on intact ABA biosynthesis for repressing some target genes, but in the absence of intact ABA biosynthesis, auxin regulates transcription of a largely distinct set of genes. In addition, our finding that there was overlap in genes that are auxin-responsive between *aba2* and wild type is consistent with our phenotypic data in that the ABA biosynthesis mutants were less responsive to auxin than wild type but still responded to auxin. Therefore, our phenotypic data and our transcript profiling data support a model whereby some parts of auxin-mediated regulation of the hypocotyl is ABA-dependent, but not all auxin-mediated regulation of the hypocotyl requires ABA.

From the ABA-treated samples, we identified a subset of 24 auxin-responsive transcripts that both require ABA2 for auxin-responsiveness and directly respond to ABA treatment (Figure 6A). This suggests that some but not all transcripts that require intact ABA biosynthesis for auxin-responsiveness are regulated by altered ABA levels in response to auxin.

**Figure 6.**
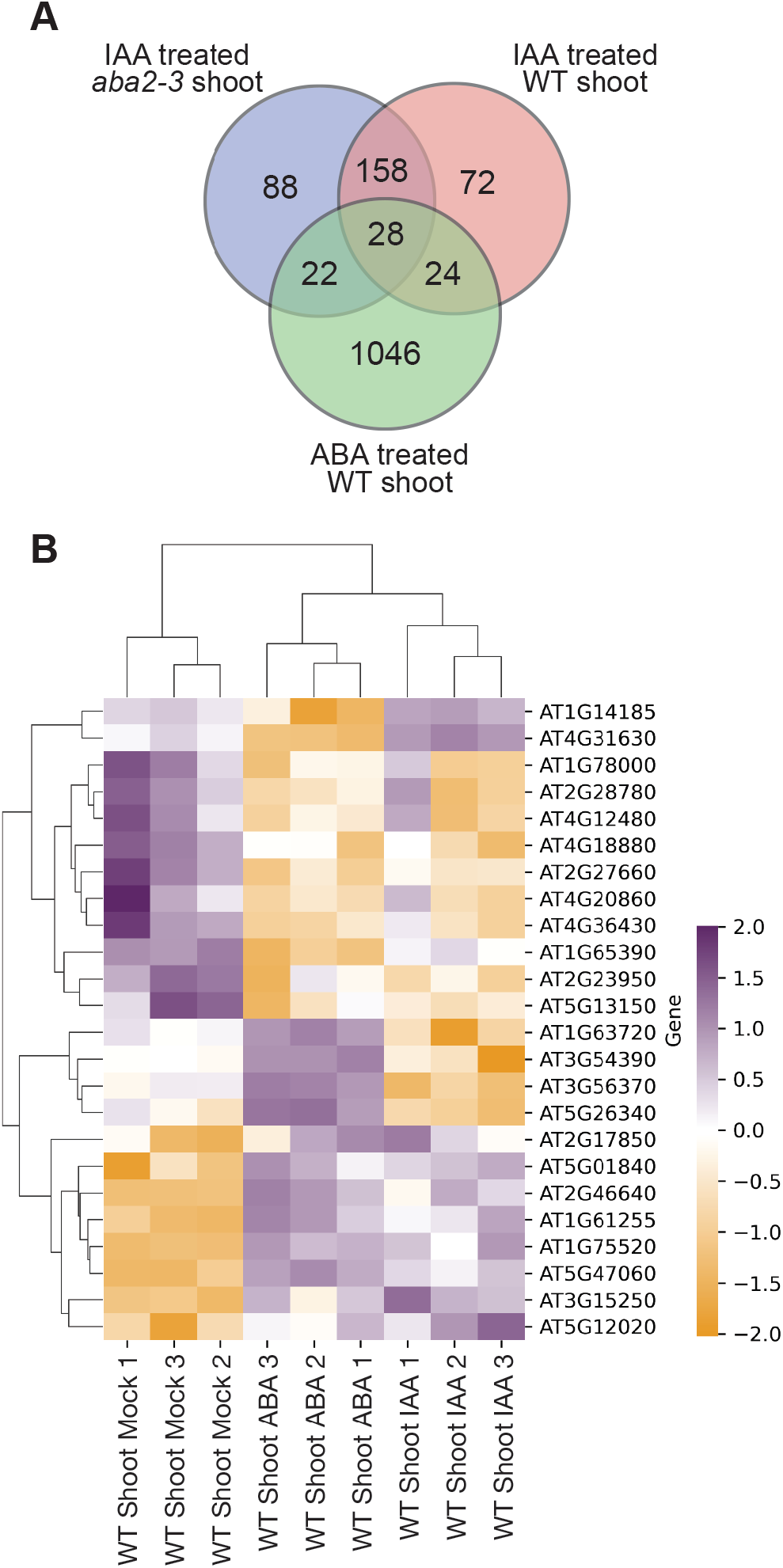
RNA-seq evaluating ABA-responsive genes identifies genes that require ABA biosynthesis for auxin response and are ABA responsive. (A) Venn diagram showing overlap between the datasets of genes that are differentially expressed (log fold change ≥ |1.0|; FDR ≤ 0.05) in shoot tissues. (B) Hierarchical clustering of the 24 genes in wild type (Wt; Col-0) shoots that both require ABA2 to respond to auxin treatment and respond directly to ABA treatment.

In addition, we examined transcript accumulation from genes that have been directly or indirectly associated with promoting or inhibiting hypocotyl elongation (Supplemental File 3), including genes involved in the biosynthesis, catabolism, or signaling of plant hormones implicated in regulating hypocotyl elongation such as GA and brassinosteroids, genes implicated in cell wall remodeling or acidification, and other genes that have been experimentally implicated in regulating hypocotyl elongation (24, 36, 37). Notably, several genes with known roles in hypocotyl elongation regulation display ABA-dependent regulation in dark-grown hypocotyls (Table 1). In particular, auxin upregulation of *GA20OX2*, a gene encoding a rate-limiting step in gibberellin biosynthesis, did not occur in the shoots of *aba2-3* mutants. GA biosynthesis is necessary for auxin to regulate hypocotyl elongation (24); our data suggest that ABA affects auxin regulation of this critical step.

**Table 1.**
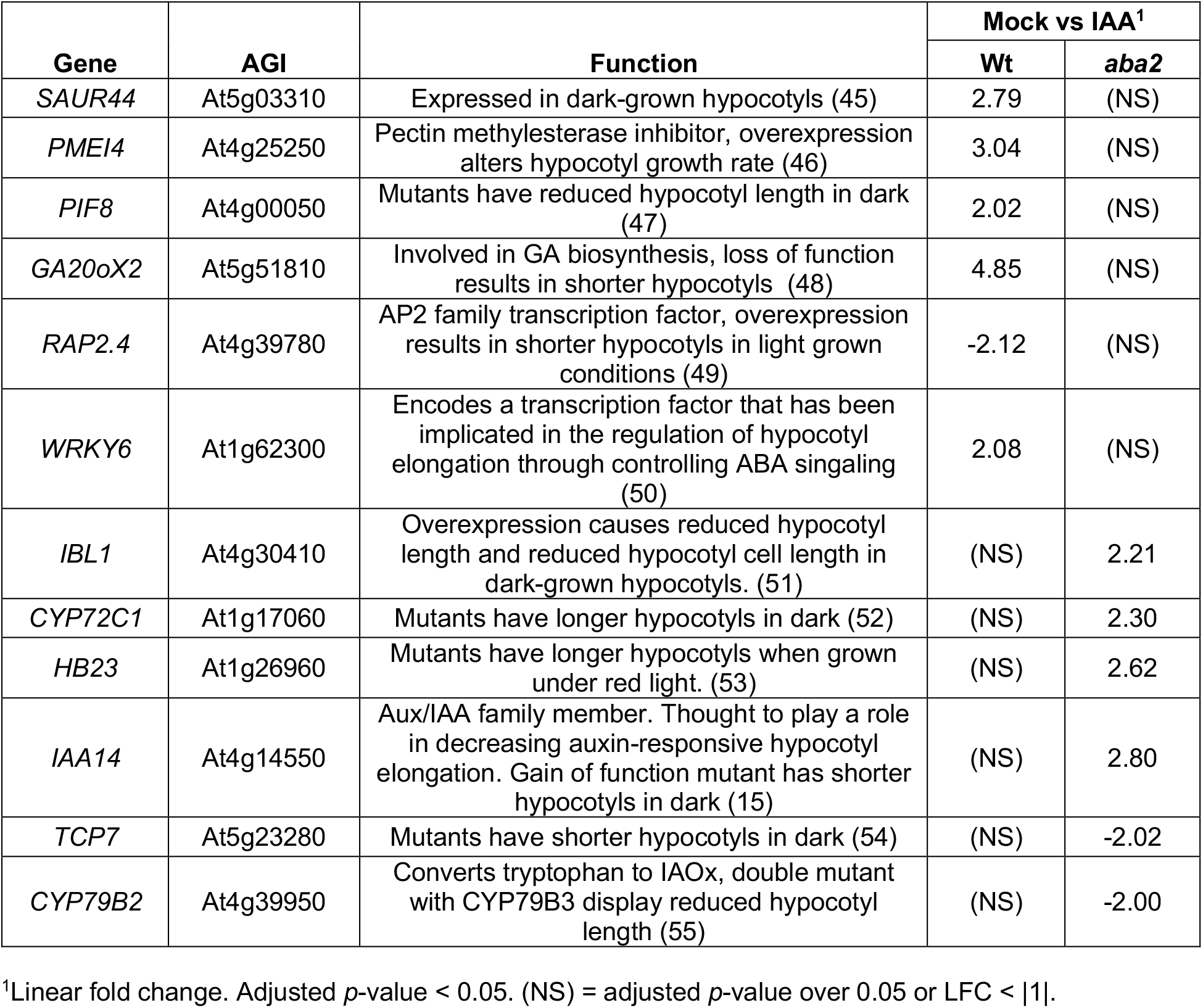
Shoot-specific ABA2-dependent auxin-responsive genes

We found multiple differences in auxin-responsive transcript accumulation from genes with known roles in hypocotyl elongation between wild type and *aba2*. No single difference is likely to explain why ABA biosynthesis mutants are resistant to the inhibitory effects of auxin on dark-grown hypocotyl elongation; however, taken together, the differential regulation of these genes by auxin in the *aba2* background may in part explain why ABA biosynthesis mutants display resistance to the effects of auxin on hypocotyl elongation. The complexity of these interactions illustrates the enigmatic connections between these pathways.

## Discussion

Hormone interactions in plants are complex. Our data shed light on auxin-ABA interactions in hypocotyl elongation. Because hypocotyl elongation is driven solely by cell expansion, dissecting hormone signaling interactions in this tissue is somewhat simpler compared to understanding hormone interactions in developmental events requiring a combination of cell division and cell expansion. However, even in a ‘simple’ system like the hypocotyl, we were unable to find a clear, single mechanism that explains the mild auxin resistance that the ABA biosynthesis mutants had in our hypocotyl elongation assays. Nonetheless, we were able to identify some genes that have previously been shown to be involved in hypocotyl elongation and require intact ABA biosynthesis for auxin responsiveness, and the additive impact of the altered regulation of these genes in the absence of normal endogenous ABA levels may in part explain the observed auxin resistance of the ABA biosynthesis mutants in hypocotyl elongation assays.

By comparing the genes regulated by auxin in the shoot of wild type seedlings to the shoot of *aba2-3* seedlings, we found striking differences in genes regulated by auxin when ABA biosynthesis was disrupted (Figure 5A). This shows that while the total number of genes that are auxin responsive in the shoot of wild type and *aba2-3* plants is largely the same (Figure 4B), the set of genes that are auxin responsive are very different in *aba2-3*. Interestingly, we found less overlap between genes downregulated by auxin in *aba2-3* and wild type shoots than those that were upregulated (Figure 5C). This suggests that disruption of ABA biosynthesis has a larger impact on the genes that auxin is able to downregulate than those that auxin upregulates. Additionally, we found that in *aba2-3* over a hundred genes that were not auxin responsive in wild type shoots became auxin responsive (Figure 5A). This suggests that not only does auxin rely on intact ABA biosynthesis to properly regulate a subset of auxin-responsive genes, but disruption of ABA biosynthesis also results in auxin gaining the capacity to regulate additional genes that are otherwise not auxin responsive.

In contrast to our understanding of auxin-regulated activation of gene transcription, auxin-triggered gene repression is not well understood. Our data provide some clues to how auxin might induce the repression of gene targets; in the case of genes repressed by auxin treatment of dark-grown hypocotyls, ABA biosynthesis is required for repression of at least a subset of targets. This finding raises the possibility that auxin-induced downregulation of genes may be through secondary pathways, such as ABA, in other tissues as well.

We found that auxin and ABA transcriptional responses in dark-grown roots and shoots are largely distinct (Figure 4). This suggests that while there are some interactions between these two hormones in regulating aspects of the shoot, specifically hypocotyl elongation, these two hormones largely regulate the transcript accumulation of non-overlapping sets of genes in the root and the shoot. This is consistent with our finding that ABA biosynthesis mutants show a mild resistance to the suppressive effects of auxin on dark-grown hypocotyl elongation and are not completely resistant to auxin like some lines containing mutations in genes encoding for core parts of the auxin signaling pathway. In addition, we found that ABA and IAA treatment often had opposing effects on the transcript accumulation of genes significantly differentially regulated in the shoot or the root by IAA and ABA treatment in wild type shoots. This suggests that while interactions between these two phytohormones play some role in regulating hypocotyl elongation, they have largely distinct impacts on gene expression. Future research into the drivers of these distinctions will be informative. Could these differences be because of expression patterns of the components of the auxin response system? Or is it possible that these context-dependent differences are due to chromatin landscape differences in different tissues? Delving into these distinctions will shed light on to how auxin plays distinct roles in different contexts.

Whereas our phenotypic analyses and transcript profiling experiments support a model whereby parts of auxin-mediated regulation of hypocotyl elongation require intact ABA biosynthesis, interactions between auxin and ABA in the regulation of this process are likely far more complex than what we have proposed. Indeed, examining the 72 genes shoot-specific genes that depend on ABA biosynthesis for auxin regulation revealed many genes involved in aspects of other phytohormone pathways, including those associated with aspects of the gibberellin, brassinosteroid, salicylic acid, jasmonate, cytokinin, and ethylene pathways. Thus, regulation of ‘simple’ hypocotyl elongation is extraordinarily complex. Our finding that auxin relies on intact ABA biosynthesis to exert its full regulatory capacity on hypocotyl elongation provides one piece of information that brings us closer to disentangling the complexities of phytohormone-regulated hypocotyl elongation.

In this study, we found that auxin requires ABA for its full inhibitory effects on dark-grown hypocotyl elongation or promotion of light-grown hypocotyls. Notably, the hypocotyl in *Arabidopsis* elongates solely via cell expansion whereas the root elongates through a combination of cell expansion and division (38). Whether auxin requires intact ABA biosynthesis to regulate other growth processes that are primarily based on cell expansion is an interesting open question. Finally, our finding that auxin requires ABA biosynthesis in order to fully regulate hypocotyl elongation suggests the possibility that stress inputs are downstream modulators of auxin-regulated growth in hypocotyl tissues, which has implications for seedling establishment.

## Materials and Methods

### Plant growth

All Arabidopsis lines with the exception of *abi1-1* and *aba2-1* were in the Col-0 background, which was used as the wild type (Wt). The *abi1-1* and *abi2-1* lines are in the L*er* background, and for assays using these lines L*er* was used as the wild type (Wt). Before plating, seeds were surface sterilized as described in (39) and then resuspended in 0.1% agar. The seeds were then stratified for 2 days at 4°C. After stratification, the seeds were plated on plant nutrient (PN) media (40) containing 0.6% agar and supplemented with 0.5% sucrose (PNS). Seedlings were grown in continuous light at 22°C.

### Phenotypic assays

For dark-grown hypocotyl elongation assays, stratified seeds were plated on PNS supplemented with hormones or the equivalent amount of ethanol for mock treatment and then exposed to continuous light for 24 hours filtered through a thin yellow long-pass plexiglass filter to help reduce breakdown of the added hormones (41). Next, the plates were wrapped in three layers of aluminum foil to ensure complete blocking of light. The seedlings were then allowed to grow for five days before hypocotyls were measured.

For light-grown hypocotyl elongation assays, stratified seeds were plated on PNS supplemented with picloram or the equivalent amount of DMSO for mock treatment. The seedlings then grew for six days under continuous light filtered through a thin yellow long-pass plexiglass filter to help reduce breakdown of the added hormones (41). The seedlings were then imaged, and the hypocotyls were measured using Image J software.

For root elongation assays, stratified seeds were plated on PNS supplemented with hormones or the equivalent amount of ethanol for mock treatment. The seedlings were grown under continuous light filtered through a thin yellow long-pass plexiglass filter to help reduce breakdown of added hormones (41). After eight days of growth, roots were measured. For statistical analysis of data acquired for the various phenotypic assays, first all groups were compared using a one-way ANOVA test to test for significant differences between the groups. In addition, pair-wise comparisons between groups using the Tukey HSD test were carried out to identify statistically significant differences between specific groups. A *p*-value of less than 0.01 was used as the cutoff for statistical significance.

### EMS mutagenesis and HR mutant isolation

Ethyl methane-sulfonate (EMS) Col-0 seeds were mutagenized as described in (42). The M_2_ seeds were then surface sterilized (39) and stratified for two days and then plated on PNS containing 30⌈M IBA. The seeds were then incubated for 24 hours under continuous light filtered through a thin yellow long-pass plexiglass filter to help reduce breakdown of the added hormones (41). The plates were then wrapped three times in aluminum foil and incubated in darkness for four days. At this time, the seedlings were screened for those exhibiting resistance to the inhibitory effects of auxin on dark-grown hypocotyl elongation. Resistant seedlings were transferred to PNS plates using sterile technique. Once large enough, the seedlings were transferred to soil. The M_3_ progeny from the auxin-resistant isolates were then re-tested for auxin resistance.

### Whole Genome Sequencing

EMS-induced mutations were identified using a whole genome sequencing of bulk segregants approach as described in (18). Briefly, seedlings isolated from the hypocotyl resistance screen were back crossed to Col-0. The F_2_ generation from this cross was then used to re-isolate seedlings exhibiting resistance to auxin in dark-grown hypocotyl elongation assays. The resultant F_3_ seed from the re-isolated seedlings was then grown and DNA isolated from the tissue of six F_3_ lines was pooled together and used for whole genome sequencing. The sequencing data was then analyzed to identify EMS-generated mutations.

### Vector construction and Arabidopsis transformation

AAO3 was amplified from *Arabidopsis thaliana* (Columbia) genomic DNA using Pfx platinum (Life Technologies) polymerase and the primers 5’-caccATGGATTTGGAGTTTGCAGTTAATGG-3’ and 5’-GTTGCTTACTTGCTTTGCCTTTATTGTC-3’. The generated PCR product was then cloned into pENTR/D-TOPO (Life Technologies) resulting in pENTR-AAO3. The pENTR-AAO3 vector was sequenced to confirm accurate cloning of AAO3. The sequenced pENTR-AAO3 vector was then recombined into the plant expression vectors pEG100 and pEG201 (43), both of which were confirmed by sequencing. The pEG100 and pEG201 vectors were then transformed into *Agrobacterium tumefaciens* strain GV3101 (44). The *Agrobacterium tumefaciens* strains carrying the pEG100 and pEG201 vectors were then used to transform Columbia and HR12 plants via the floral dip method. From the dipped plants, T_1_ seed was harvested, surface sterilized, and then plated on PN media supplemented with 10 ⌈g/mL BASTA (40). Subsequent generations were tested to identify lines homozygous for the transgene, which were then used for downstream experiments.

### RNA isolation and RNA-seq experiment

For the RNA-seq experiment, Col-0 (Wt) and *aba2-3* seeds were surface sterilized (39), resuspended in agar, and then incubated for two days at 4°C. The seeds were then plated on PNS plates and incubated for 24 hours under continuous light. The plates were then wrapped in three sheets of aluminum foil and incubated for four days. Next, the dark-grown plants were transferred to liquid PNS containing either 10⌈M IAA, 10⌈M ABA, or the equivalent amount of ethanol for the mock treatment in a dark room with only a green safe-light for illumination. Each treatment was repeated in triplicate for both lines. After incubation for two hours, the seedlings were removed from the liquid PNS, cut and separated at the root shoot junction, and the isolated tissue was then immediately frozen in liquid nitrogen. Total RNA was isolated from each tissue for each respective treatment using the RNeasy Plant Mini Kit from Qiagen according to manufacturer’s instructions. Samples were then prepared for sequencing using the SMARTer cDNA synthesis kit (Clontech) according to manufacturer’s instructions. The samples were sequenced across three (1×50bp) lanes using an Illumina HiSeq 3000. The reads were then demultiplexed and aligned to the TAIR 10 Ensemble Release 23 assembly using STAR. Gene counts were derived from the number of uniquely aligned unambiguous reads by Subread:featureCount.

Gene counts were normalized using the R/Bioconductor package EdgeR, and TMM normalization size factors were calculated to adjust values due to differences in library size. Ribosomal genes and genes not expressed in any sample greater than 2 counts-per-million were not considered for further analysis. The calculated TMM size factors and the matrix of counts were then imported into the R/Bioconductor Limma package, and these values were used to make a Spearman correlation matrix (Figure S1A) and multi-dimensional scaling plot (Figure S1B). The weighted likelihoods based on observed mean-variance relationship of every gene for every sample were calculated using the voomWithQualityWeights function. The residual standard deviation of every gene to the gene’s average log-count was plotted to assess gene performance, and these values were used to fit a trendline of the residuals (Figure S1C). A generalized linear model was then created to test for gene level differential expression, and results for genes with a Benjamini-Hochberg flase-discovery rate adjusted p-value of greater than or equal to 0.05 were filtered out.

### Gene ontology analysis

Gene ontology analysis was carried out using panther.db (25). Genes were identified by searching for statistically overrepresented genes within the different lists of differentially expressed for the various comparisons. The specific analysis used was the statistical overrepresentation test using the GO biological process complete annotation set.

## Data availability

The RNA-seq data discussed in this publication have been deposited in NCBI’s Gene Expression Omnibus and are accessible through GEO Series accession number GEO: GSE169302.

## Acknowledgments

We thank Nicholas Morffy, Joe Cammarata, Hongwei Jing and Sunita Pathak for helpful comments. This research was supported by the William H. Danforth Plant Science Fellowship Program (to R.J.E), the National Science Foundation Center for Engineering Mechanobiology (CMMI-1548571 to L.C.S.), and the National Institutes of Health (R35 GM136338 to L.C.S.). We thank the Genome Technology Access Center in the Department of Genetics at Washington University School of Medicine for help with RNA-seq analyses.

---

**Supplemental Figure 1.**
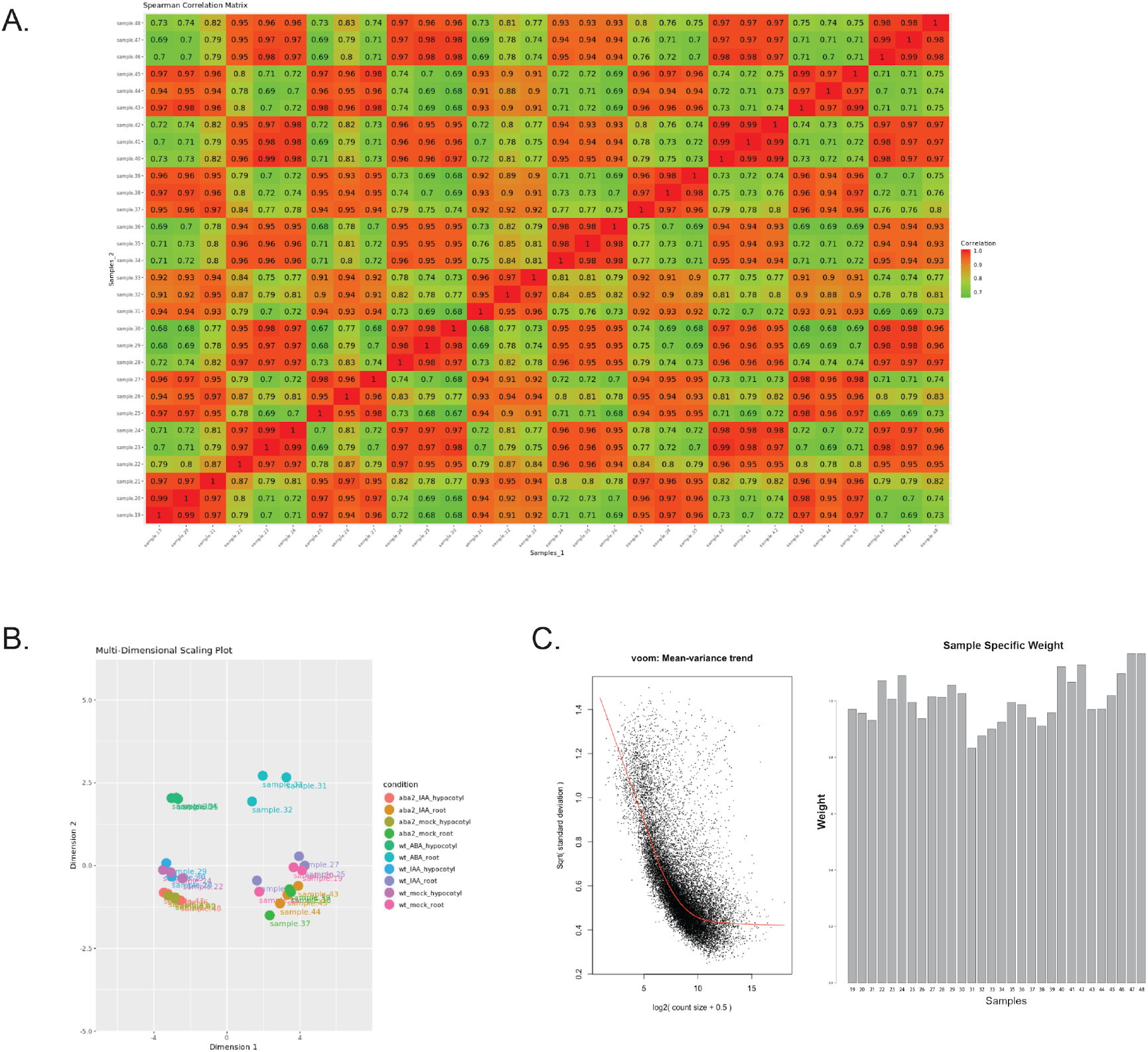
RNA-seq Sample Quality Assessment. (A) Spearman correlation values for all genes with a count-per-million value of greater than 1 in at least 3 samples. (B) Grouping of various samples using a multi-dimensional scaling plot of the leading log fold changes for the various samples. (C) The voom:Mean-variance trend plot (left) shows a scatter plot of empirically derived fitted and trended mean-variance relationship across all genes. The sample-specific weights plot (right) shows the sample specific weights for each sample for the statistical model fitting and differential transcript accumulation analysis.

## References

1. M. Vanstraelen, E. Benková, Hormonal interactions in the regulation of plant development. Annu Rev Cell Dev Biol 28, 463–487 (2012).

2. C. Perrot-Rechenmann, Cellular responses to auxin: division versus expansion. Cold Spring Harb Perspect Biol 2, a001446 (2010).

3. T. Yoshida et al., The Role of Abscisic Acid Signaling in Maintaining the Metabolic Balance Required for Arabidopsis Growth under Nonstress Conditions. Plant Cell 31, 84–105 (2019).

4. R. J. Emenecker, L. C. Strader, Auxin-Abscisic Acid Interactions in Plant Growth and Development. Biomolecules 10 (2020).

5. X. Liu et al., Auxin controls seed dormancy through stimulation of abscisic acid signaling by inducing ARF-mediated ABI3 activation in Arabidopsis. Proc Natl Acad Sci U S A 110, 15485–15490 (2013).

6. J. M. Thole, E. R. Beisner, J. Liu, S. V. Venkova, L. C. Strader, Abscisic acid regulates root elongation through the activities of auxin and ethylene in Arabidopsis thaliana. G3 (Bethesda) 4, 1259–1274 (2014).

7. C. Belin, C. Megies, E. Hauserova, L. Lopez-Molina, Abscisic acid represses growth of the Arabidopsis embryonic axis after germination by enhancing auxin signaling. Plant Cell 21, 2253–2268 (2009).

8. M. Monroe-Augustus, B. K. Zolman, B. Bartel, IBR5, a dual-specificity phosphatase-like protein modulating auxin and abscisic acid responsiveness in Arabidopsis. Plant Cell 15, 2979–2991 (2003).

9. R. Lorrai et al., Abscisic acid inhibits hypocotyl elongation acting on gibberellins, DELLA proteins and auxin. AoB Plants 10, ply061 (2018).

10. L. C. Strader, M. Monroe-Augustus, B. Bartel, The IBR5 phosphatase promotes Arabidopsis auxin responses through a novel mechanism distinct from TIR1-mediated repressor degradation. BMC Plant Biol 8, 41 (2008).

11. C. Lu et al., Abscisic Acid Regulates Auxin Distribution to Mediate Maize Lateral Root Development Under Salt Stress. Front Plant Sci 10, 716 (2019).

12. L. Xing, Y. Zhao, J. Gao, C. Xiang, J. K. Zhu, The ABA receptor PYL9 together with PYL8 plays an important role in regulating lateral root growth. Sci Rep 6, 27177 (2016).

13. R. Shin et al., The Arabidopsis transcription factor MYB77 modulates auxin signal transduction. Plant Cell 19, 2440–2453 (2007).

14. D. Shkolnik-Inbar, D. Bar-Zvi, ABI4 mediates abscisic acid and cytokinin inhibition of lateral root formation by reducing polar auxin transport in Arabidopsis. Plant Cell 22, 3560–3573 (2010).

15. M. A. Rinaldi, J. Liu, T. A. Enders, B. Bartel, L. C. Strader, A gain-of-function mutation in IAA16 confers reduced responses to auxin and abscisic acid and impedes plant growth and fertility. Plant Mol Biol 79, 359–373 (2012).

16. L. C. Strader et al., Multiple facets of Arabidopsis seedling development require indole-3-butyric acid-derived auxin. Plant Cell 23, 984–999 (2011).

17. M. Seo et al., The Arabidopsis aldehyde oxidase 3 (AAO3) gene product catalyzes the final step in abscisic acid biosynthesis in leaves. Proc Natl Acad Sci U S A 97, 12908–12913 (2000).

18. J. M. Thole, L. C. Strader, Next-generation sequencing as a tool to quickly identify causative EMS-generated mutations. Plant Signal Behav 10, e1000167 (2015).

19. L. M. Ng, K. Melcher, B. T. Teh, H. E. Xu, Abscisic acid perception and signaling: structural mechanisms and applications. Acta Pharmacol Sin 35, 567–584 (2014).

20. W. Yang, W. Zhang, X. Wang, Post-translational control of ABA signalling: the roles of protein phosphorylation and ubiquitination. Plant Biotechnol J 15, 4–14 (2017).

21. B. Brandt et al., Reconstitution of abscisic acid activation of SLAC1 anion channel by CPK6 and OST1 kinases and branched ABI1 PP2C phosphatase action. Proc Natl Acad Sci U S A 109, 10593–10598 (2012).

22. Y. Ma et al., Regulators of PP2C phosphatase activity function as abscisic acid sensors. Science 324, 1064–1068 (2009).

23. Y. Hayashi, K. Takahashi, S. Inoue, T. Kinoshita, Abscisic acid suppresses hypocotyl elongation by dephosphorylating plasma membrane H(+)-ATPase in Arabidopsis thaliana. Plant Cell Physiol 55, 845–853 (2014).

24. E. J. Chapman et al., Hypocotyl transcriptome reveals auxin regulation of growth-promoting genes through GA-dependent and -independent pathways. PLoS One 7, e36210 (2012).

25. H. Mi, A. Muruganujan, P. D. Thomas, PANTHER in 2013: modeling the evolution of gene function, and other gene attributes, in the context of phylogenetic trees. Nucleic Acids Res 41, D377–386 (2013).

26. S. K. Powers et al., Nucleo-cytoplasmic Partitioning of ARF Proteins Controls Auxin Responses in Arabidopsis thaliana. Mol Cell 76, 177–190 e175 (2019).

27. N. A. Omelyanchuk et al., Auxin regulates functional gene groups in a fold-change-specific manner in Arabidopsis thaliana roots. Sci Rep 7, 2489 (2017).

28. S. F. Zia et al., Direct comparison of Arabidopsis gene expression reveals different responses to melatonin versus auxin. BMC Plant Biol 19, 567 (2019).

29. T. T. Yuan, H. H. Xu, Q. Zhang, L. Y. Zhang, Y. T. Lu, The COP1 Target SHI-RELATED SEQUENCE5 Directly Activates Photomorphogenesis-Promoting Genes. Plant Cell 30, 2368–2382 (2018).

30. G. Qin et al., An indole-3-acetic acid carboxyl methyltransferase regulates Arabidopsis leaf development. Plant Cell 17, 2693–2704 (2005).

31. G. Martin et al., The photoperiodic response of hypocotyl elongation involves regulation of CDF1 and CDF5 activity. Physiol Plant 169, 480–490 (2020).

32. B. C. Kim, M. C. Soh, B. J. Kang, M. Furuya, H. G. Nam, Two dominant photomorphogenic mutations of Arabidopsis thaliana identified as suppressor mutations of hy2. Plant J 9, 441–456 (1996).

33. S. Wang, Y. Chang, J. Guo, J. G. Chen, Arabidopsis Ovate Family Protein 1 is a transcriptional repressor that suppresses cell elongation. Plant J 50, 858–872 (2007).

34. A. Frey et al., Epoxycarotenoid cleavage by NCED5 fine-tunes ABA accumulation and affects seed dormancy and drought tolerance with other NCED family members. Plant J 70, 501–512 (2012).

35. R. J. Laby, M. S. Kincaid, D. Kim, S. I. Gibson, The Arabidopsis sugar-insensitive mutants sis4 and sis5 are defective in abscisic acid synthesis and response. Plant J 23, 587–596 (2000).

36. D. S. Favero, A. Lambolez, K. Sugimoto, Molecular pathways regulating elongation of aerial plant organs: a focus on light, the circadian clock, and temperature. Plant J 10.1111/tpj.14996 (2020).

37. X. Y. Zhou, L. Song, H. W. Xue, Brassinosteroids regulate the differential growth of Arabidopsis hypocotyls through auxin signaling components IAA19 and ARF7. Mol Plant 6, 887–904 (2013).

38. E. Gendreau et al., Cellular basis of hypocotyl growth in Arabidopsis thaliana. Plant Physiol 114, 295–305 (1997).

39. R. L. Last, G. R. Fink, Tryptophan-Requiring Mutants of the Plant Arabidopsis thaliana. Science 240, 305–310 (1988).

40. G. W. Haughn, C. Somerville, Sulfonylurea-resistant mutants of Arabidopsis thaliana. Molecular and General Genetics MGG 204, 430–434 (1986).

41. T. C. Stasinopoulos, R. P. Hangarter, Preventing photochemistry in culture media by long-pass light filters alters growth of cultured tissues. Plant Physiol 93, 1365–1369 (1990).

42. J. Normanly, P. Grisafi, G. R. Fink, B. Bartel, Arabidopsis mutants resistant to the auxin effects of indole-3-acetonitrile are defective in the nitrilase encoded by the NIT1 gene. Plant Cell 9, 1781–1790 (1997).

43. K. W. Earley et al., Gateway-compatible vectors for plant functional genomics and proteomics. Plant J 45, 616–629 (2006).

44. C. Koncz, J. Schell, The promoter of TL-DNA gene 5 controls the tissue-specific expression of chimaeric genes carried by a novel type of Agrobacterium binary vector. Molecular and General Genetics MGG 204, 383–396 (1986).

45. N. Sun et al., Arabidopsis SAURs are critical for differential light regulation of the development of various organs. Proc Natl Acad Sci U S A 113, 6071–6076 (2016).

46. S. Pelletier et al., A role for pectin de-methylesterification in a developmentally regulated growth acceleration in dark-grown Arabidopsis hypocotyls. New Phytol 188, 726–739 (2010).

47. J. Oh, E. Park, K. Song, G. Bae, G. Choi, PHYTOCHROME INTERACTING FACTOR8 Inhibits Phytochrome A-Mediated Far-Red Light Responses in Arabidopsis. Plant Cell 32, 186–205 (2020).

48. A. R. Plackett et al., Analysis of the developmental roles of the Arabidopsis gibberellin 20-oxidases demonstrates that GA20ox1, -2, and -3 are the dominant paralogs. Plant Cell 24, 941–960 (2012).

49. R. C. Lin, H. J. Park, H. Y. Wang, Role of Arabidopsis RAP2.4 in regulating light-and ethylene-mediated developmental processes and drought stress tolerance. Mol Plant 1, 42–57 (2008).

50. R. Lorrai et al., Genome-wide RNA-seq analysis indicates that the DAG1 transcription factor promotes hypocotyl elongation acting on ABA, ethylene and auxin signaling. Sci Rep 8, 15895 (2018).

51. M. K. Zhiponova et al., Helix-loop-helix/basic helix-loop-helix transcription factor network represses cell elongation in Arabidopsis through an apparent incoherent feed-forward loop. Proc Natl Acad Sci U S A 111, 2824–2829 (2014).

52. N. Takahashi et al., shk1-D, a dwarf Arabidopsis mutant caused by activation of the CYP72C1 gene, has altered brassinosteroid levels. Plant J 42, 13–22 (2005).

53. H. Choi et al., The homeodomain-leucine zipper ATHB23, a phytochrome B-interacting protein, is important for phytochrome B-mediated red light signaling. Physiol Plant 150, 308–320 (2014).

54. G. Zhang et al., TCP7 functions redundantly with several Class I TCPs and regulates endoreplication in Arabidopsis. J Integr Plant Biol 61, 1151–1170 (2019).

55. Y. Zhao et al., Trp-dependent auxin biosynthesis in Arabidopsis: involvement of cytochrome P450s CYP79B2 and CYP79B3. Genes Dev 16, 3100–3112 (2002).

